# Developing a 3- to 6-state EEG-based brain-computer interface for a robotic manipulator control

**DOI:** 10.1101/171025

**Authors:** Yuriy Mishchenko, Murat Kaya, Erkan Ozbay, Hilmi Yanar

## Abstract

Recent developments in BCI techniques have demonstrated high-performance control of robotic prosthetic systems primarily via invasive methods. In this work we develop an electroencephalography (EEG) based noninvasive BCI system that can be used for a similar, albeit lower-speed robotic control, and a signal processing system for detecting user’s mental intent from EEG data based on up to 6-state motor-imagery BCI communication paradigm. We examine the performance of that system on experimental data collected from 12 healthy participants and analyzed offline. We show that our EEG BCI system can correctly identify different motor imageries in EEG data with high accuracy: 3 out of 12 participants achieved accuracy of 6-state communication in 80-90% range, while 2 participants could not achieve a satisfactory accuracy. We further implement an online BCI system for control of a virtual 3 degree-of-freedom prosthetic manipulator and test it with our 3 best participants. The participants’ ability to control the BCI is quantified by using the percentage of successfully completed BCI tasks, the time required to complete a task, and the error rate. 2 participants were able to successfully complete 100% of the test tasks, demonstrating on average the error rate of 80% and requiring 5-10 seconds to execute a manipulator move. 1 participant failed to demonstrate a satisfactory performance in online trials. Our results lay a foundation for further development of EEG BCI-based robotic assistive systems and demonstrate that EEG-based BCI may be feasible for robotic control by paralyzed and immobilized individuals.

## I. Introduction

Patients paralyzed due to trauma or accident or immobilized due to ongoing medical treatment suffer a significant loss of motor and communication ability. Recent advances in neural prosthetics have the potential to allow such patients to regain control of a multitude of motor and communication functions [1]–[3]. Targeted muscle re-innervations [4] and myoelectric control using residual muscle activity offer exciting possibilities in this respect [5]–[7]. Another option is provided by brain-computer interfaces (BCI) that directly translate neural activity in the cortex into a control signal for external devices [8]–[13]. Recent advances in BCI research include neural control of robotic manipulators realized in monkeys [14]–[17], nonhuman primates [18]–[20], and virtual 2D/3D control by paralyzed individuals [21]–[25]. [24] reported results for a tetraplegic human patient implanted with a 96-microelectrode array in the primary motor cortex area. The patient was able to achieve, in the course of 57 consecutive sessions over 9 months, a high-performance BCI control of 2D cursor movements that could be used to read e-mail, operate external devices, or transport objects by means of a robotic manipulator. More recent results demonstrated the ability of BCI for high-performance control of 3D reach and grasp by a tetraplegic patient [26]. In recent work [27], a tetraplegic patient has been shown to be able to learn the control of a 7 dof robotic manipulator arm using BCI. In that regard, intracranial invasive BCI have shown significant potential for control of robotic prosthetic technology with a high degree of efficiency and accuracy [19], [26], [27]. Despite these impressive achievements, intracranial BCI represent significant risks to their users associated with the necessarily of invasive brain surgery. Thus, exploring the potential for a high-performance control of brain-computer interfacing technologies using only noninvasive brain activity imaging means is of great interest.

Information about users’ motor intent has been shown in the literature to exist in cortical activity in a variety of frequency bands and spatial scales from local field potentials (LFP), to electrocorticography (ECoG), functional Magnetic Resonance Imaging (fMRI), magnetoencephalography (MEG) and electroencephalography (EEG) scales [28]–[32]. In that relation, EEG represents a point of special interest given the ease with which EEG data can be acquired, the maturity of the technology, the portability, versatility and relatively low cost of modern EEG acquisition devices as contrasted with invasive and other noninvasive brain activity imaging techniques. In the past, EEG-based BCIs have been used for a diverse set of applications. The first EEG BCI developed by Vidal [33] in 1973 enabled the users to control the direction of cursor movement using eye gaze. After that pioneering work, Wolpaw and McFarland with coworkers developed an EEG

BCI based on sensorimotor rhythm modulation for 1D, 2D and 3D computer cursor control [25], [34], [35]. EEG based BCIs have been used not only for computer cursor control but also for control of assistive devices such as robotic manipulators and motorized wheelchairs [36]–[38]. In [36] an EEG BCI-based system for control of the direction of motion of a wheelchair had been described. The RIKEN BSI-TOYOTA Collaboration Center has succeeded in providing a real-time control of a wheelchair with 95% accuracy by using an EEG BCI. Pfurtscheller et al. have described a BCI neurostimulation system that used EEG signals to control a functional electrical stimulation of a tetraplegic patient’s hand and allowed the patient to grasp a cylinder using her hand and this setup [37]. Chae et al. have applied EEG BCI to achieve an asynchronous control of navigation of a humanoid robot by the users. They have performed a series of experiments organized in the format of offline training, online feedback testing, and real-time control sessions, and reported a success rate of 93.3% accuracy in one of five participants, who use the BCI to controlled the robot navigating in an indoor maze [39]. In another recent application of EEG BCI, Sankai developed a robotic exoskeleton suit named HAL (Hybrid Assistive Limb), aimed for medical and human-enhancement applications [40]. Kwak et al. have developed a similar asynchronous EEG BCI-based lower limb exoskeleton system controlled using steady-state visual evoked potentials [41]. [42] describes a hybrid hand exoskeleton for restoration of independent daily activities of six paraplegic individuals using EEG BCI.

In this paper, we develop a noninvasive EEG-based BCI that can be applied in the future towards a high performance control of a robotic manipulator envisioned as a part of an assistive robotic complex for paralyzed or immobilized patients, similar to that discussed in [19], [26], [27]. The paper is organized as follows. Section II describes the process of EEG data collection, pre-processing, and analysis both for our offline and online BCI applications. In Section III, the ability of the developed system to discriminate different mental imagery states is examined on 12 participants using offline analysis of collected EEG data. Certain design choices of the BCI’s data processing system including that of EEG data representation, detector window optimization, reference voltage choice, frequency filter, etc. are investigated. The results of interactive application of BCI for online control of a 3D robotic arm simulated on a computer screen by 3 participants are also presented. In Section IV, we discuss achieved results and compare them with the literature. Achieved performance and venues for potential improvement as well as the perspectives of the BCI’s utilization in assistive robotic settings are discussed. Conclusions are offered in Section V.

## II. Materials and Methods

### A. Data acquisition and experimental procedures

The EEG data was acquired using the medical grade EEG-1200 EEG recording system with JE-921A acquisition box (Nihon Kohden, Japan). A standard EEG cap (Electro-Cap International, USA) with 19 electrodes in 10-20 configuration was used for all experiments. Before each experiment, participants had their head prepared for recording EEG data by cleaning the surface of skin with alcohol solution and combing hair around the locations of electrode-sites. After preparation, the EEG cap was carefully placed onto participant’s head; for that, the distances to nasion, inion and preaurical points from Cz electrode were measured and the correct placement of the cap was ensured. Once the correct position of the EEG cap had been ensured, the electrodes were filled with electro-gel (Electro-Gel, Elector-Cap International, USA; Elefix Paste for EEG Z-401CE, Nihon Kohden, USA) while simultaneously controlling the electrode impedances in the impedance-check mode of EEG 1200 Neurofax program. After achieving all the impedances at or below 10 kOhm with the impedance imbalance at or below 5 kOhm, the EEG recordings preparation was complete.

The EEG experiments were conducted in two formats - offline format and online format. In the offline format, the EEG data was first collected following the procedures described below, and the analysis of the EEG data was performed offline after the experiment. In the online format, the participants attempted to control a robot manipulator arm simulated on computer screen by using the BCI system in online, interactive mode, as will be described below.

In the offline experiments, the participants were comfortably seated in a recliner chair with the EEG cap prepared and the computer screen of the original base computer part of the EEG 1200 system positioned approximately 200 cm in front of them at slightly above the eye level. The computer screen was made show the experiment’s graphical user interface (eGUI) shown in Fig. 1 and consisting of a Matlab figure with 5 icons encoding different motor imageries (left hand, right hand, left leg, right leg, tongue) as well as a “passive” icon represented by a circle and a gaze-fixation point in the center of the figure, see Fig. 1. Prior to the beginning of the data acquisition, the participants were instructed to remain motionless and keep their gaze fixed at the fixation point at all times during the experiment.

**Fig. 1.**
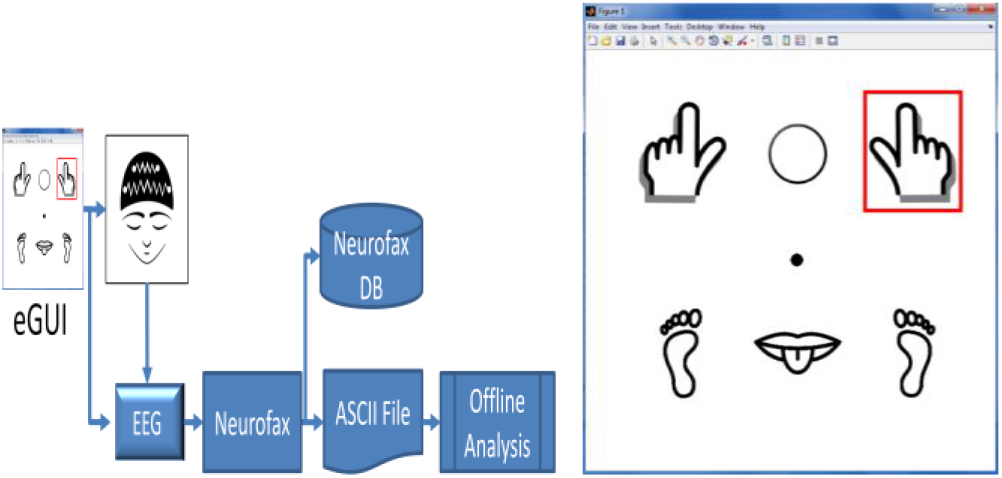
(Left) Schematic representation of the offline experiments’ procedure. First, action signals had been presented to participants by indicating one of the motor imagery icons using a red rectangle on computer screen. The EEG signal in response to the implementation of shown motor imagery was acquired by EEG-1200 system and recorded using Neurofax software. After the completion of the experiment, the recorded EEG data was saved to Neurofax database as well as exported to an ASCII file for further custom offline processing. (Right) The graphical user interface used in offline experiment consisting of a single Matlab figure with 5 icons representing left hand, right hand, left leg, right leg, and tongue motor imageries, a passive signal indicated with a circle, a fixation point in the center, and an action signal shown as a red rectangle around right hand motor imagery icons.

Each experiment began with a 2.5 minute relaxation period, after which three 15-minute sessions followed separated by 2 minute breaks, for a total duration of approximately 1 hour. During each session, participants performed approximately 300 BCI trials each consisting of one 1-second visual action signal presentation that selected a motor imagery to be implemented by means of a red rectangle shown around the corresponding imagery’s icon in eGUI, Fig. 1. The motor imageries were selected randomly following a pseudorandom sequence generated in the beginning of the experiment using a random number generator. The participants were instructed to implement the shown imagery for one time during the period in which the selection frame was visible. A “passive” action signal was shown using a circle icon. During the passive signal, the participants were instructed to remain passive and to not intentionally respond in any way. A pause of duration varying from 1.5 to 2.5 seconds followed the presentation of each action signal, thus concluding the trial. Each trial took on average 3 seconds to complete.

The EEG signals from 19 standard EEG electrodes plus two A1-A2 ground electrodes were recorded during the entire duration of the experiment using Neurofax software (Nihon Kohden, Japan). The recording settings used were the sampling rate of 200 Hz, the low frequency cut-off of 0.53 Hz, and the high frequency cut off 70 Hz. A custom montage including all the electrodes with system 0 Volt-reference was created in Neurofax and used in the recordings, archived data, and ASCII exported data.

The EEG data coming from EEG-1200 system needed to be additionally synchronized with the computer’s clock, according to which the times of the action signal presentations were recorded. We found that the internal timer of JE-921A acquisition box could have different offset as well as a speed different from the base computer’s system clock sufficient to destroy the observability of event-related potentials in the data. To achieve this synchronization, a 1 μV signal was forwarded from eGUI via the computer’s serial port and an Arduino-based adaptor to a bipolar input port in JE-921A acquisition box (namely, X3), thus, marking the onset time of all action signals in the data received from EEG-1200 via the X3 bipolar input channel.

After the completion of each experiment, acquired EEG data was saved to Neurofax internal database as well as externally exported to an ASCII file by using the ASCII export function of Neurofax software. The exported EEG data comprised a file listing all the voltages on the recorded EEG electrodes at every point of time in a text-table format. The voltage resolution of the exported data was 0.01 μV and the sampling rate was 200 samples per second. The entire duration of the experiment was covered by the exported data. The data was then imported to Matlab for offline analysis using a custom script based on convert_nkascii2mat.m function by Timothy Ellmore available from Beauchamp at OpenWetWare.org.

In the online experiments, the participants were asked to control a 3 dof robot manipulator arm simulated on a computer screen in 3D, using our EEG BCI, Fig. 2. The robot manipulator could perform left-right, forward-backward, and hold-release motions, controlled by up to 6 mental imageries including left and right hand movement, left and right leg movement, tongue movement, and one passive imagery. The preparation and recording of the EEG signal were performed in the same way as described for the offline experiments. In order to acquire the EEG data from EEG-1200 in real time, we have developed a custom driver software using C#, given that Neurofax software did not provide options for real-time exporting raw EEG data. The said driver performed a scan of the virtual memory space of Neurofax program, identified a region in the memory associated with a ring buffer used for updating the on-screen EEG traces in Neurofax software, and then extracted that data once per each 10 msec time interval. The extracted data was then forwarded to our main BCI application in Matlab by variable sharing via the Matlab COM interface. This allowed us to acquire the raw EEG data from EEG-1200 in real time. The data acquired had the sampling rate of 200 Hz, the voltage resolution of 0.133 μV, and the dynamic range of ±100 μV. The EEG data had to be synchronized with the main computer’s system clock, as mentioned above. During the online experiments, we found that EEG-1200 not just could have different time offset and speed, but also that Neurofax software could output the data with a variable delay ranging from 300 to 700 msec. To remove this variation, we forwarded a 1-μV signal at each presentation of an action signal to the X3 bipolar input port in JE-921A acquisition box, and used that signal to precisely determine the onset time of each action signal.

**Fig. 2.**
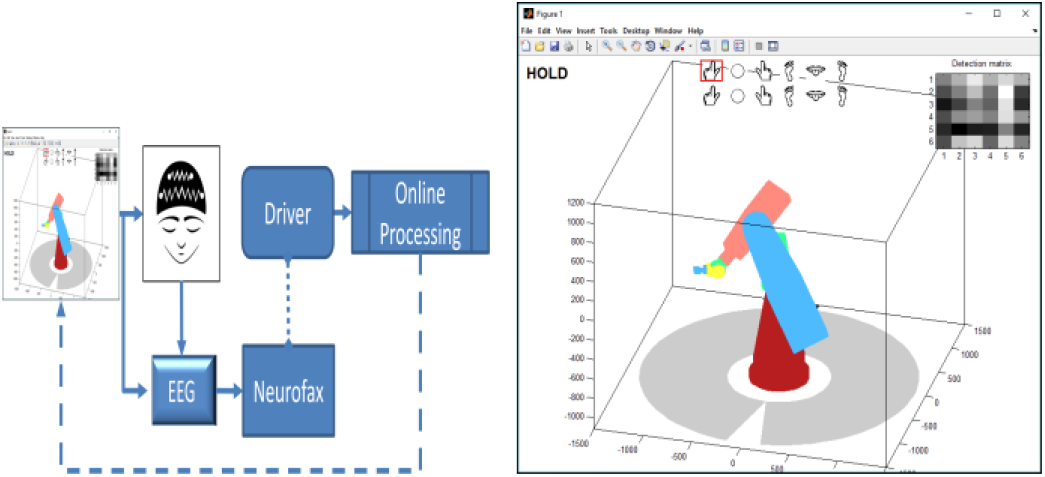
(Left) Schematic representation of the online experiments’ procedure. The participants attempted to control in real time a 3 dof robotic manipulator arm using the EEG BCI by implementing different motor imageries. The row EEG signal associated with implementation of each imagery was recorded by EEG-1200 system and imported to Matlab via custom memory-scanning driver software. The EEG data was then processed in a Matlab application and used to provide control signal for the robot arm. (Right) The online experiment’s interactive graphical user interface (iGUI) consisting of a Matlab figure modeling in 3D the motion of a robot manipulator arm.

The online experiments followed a protocol essentially similar to that used in the offline experiments. Namely, the online experiments were structured as three 15-minutes sessions separated by 2 minute breaks, with a 2.5 minute initial relaxation period. The first 15-minute session was used to train the BCI decoder. During that session, a random sequence of 1-second action signals was presented to the participants by means of the top row of motor imagery icons in the user interface, Fig. 2, selecting motor imageries that the participants needed to implement. The action signals were separated by 1.5-2.5 second break intervals for the average time per action signal of 3 seconds, exactly the same as in the offline experiments. The data collected during that session was used to construct the BCI signal decoder model as will be described following sections.

The second 15-minute session was used as a practice session. During that time, the participants used the previously trained BCI to try and assume the control of the manipulator arm in a free exploration manner. During that session, an “on” signal was shown to the participants using a red rectangle drawn over empty space in the top row of imagery icons for 1 second every 2.5-3.5 seconds. During that signal, the participants implemented the imagery of their choice in voluntary manner similar to such practiced in the training session. The BCI then attempted to recognize the imagery. The recognized imagery was used to move the simulated manipulator arm and also was shown in the second motor-imagery icons row for feedback.

The third 15-minute session was used as a test session. During that time, the participants were given instructions to execute a set of simple tasks by using the robot arm and the BCI. The tasks consisted of moving the robot arm to a specified location such as “move robot arm two steps to the left” or “move robot arm two steps to the left and two steps forward”. The participants used the same procedures to execute those tasks as described above for the practice session.

### B. Offline data analysis

The analysis of collected EEG data in the offline experiments was performed offline after the conclusion of each experiment. For that, the EEG data was first exported from Neurofax software into an ASCII file and imported to Matlab using a custom script, as described in the previous section. The EEG data thus imported to Matlab was a matrix of EEG voltage readings, *E_it_*, where *t =* 1,…, *T* indexed the different time samples and *i =* 1,…, *N_e_* indexed the EEG channels. A total of *N_e_ =* 22 data channels were imported, with 19 channels corresponding to the EEG electrodes in the standard 10-20 configuration, two channels describing the A1-A2 electrodes, and the 22^nd^ channel encoding the eGUI synchronization-signal on the X3 bipolar input, as described previously. The raw EEG data was in system 0 Volt-reference (defined by EEG-1200 manual as the average of the voltage values on C3 and C4 electrodes) and was digitally rereferenced to different voltage references including the A1-A2 average reference (defined as the average of the A1 and A2 voltages), the common average reference (defined as the average of all electrodes’ voltages), and the Laplace reference (defined as the average of the voltages on each electrode’s 4 immediate neighbors). The data in the sync-channel was used to establish the onset times of the action and to bind each action signal in eGUI to a specific onset time. Other than re-referencing and binding of the action signals, no other preprocessing was performed on raw EEG data.

The EEG data was partitioned into data frames corresponding to different presentations of action signal for further processing, by selecting the fragments of EEG data within [t_1_, t_2_]-data frames locked to the action signal onset time. The stack of such data frames could be represented as a 3-dimensional array, *E_nit_*, with the indexes *i* and *t* having the same meaning as before (except that *t* changed from 1 to *dt = t_2_ - t_1_* - the length of the frame), and the index *n* enumerating different trials or action signal presentation episodes. Generally, the first second of EEG data immediately following the onset of the action signal could be used as the data frames for decoding, which coincided with the time during which the action signal was on and the participants carried out mental imageries. However, we further optimized the data-frame selection by considering all possibilities for the frame onset, t_1_ ϵ [-0.5,1.0] sec, and end-time, t_2_ ϵ [0.0,2.ö] sec, at 0.1 second intervals. As described in Results, the decoding data frame choice of *t_1_* = 0 sec and *t_2_* = 0.85sec was found to offer the best overall performance for all subjects and all mental imageries.

After partitioning the EEG recording into data frames, the decoder for associating data frames with mental imageries was constructed using SVM or LDA machine learning algorithms. For that, first, the EEG data from each frame was converted into a feature-vector representation. We experimented with several feature representations of EEG signal including Power Spectral Density (PSD), EEG band power (EEG band), Fourier transform amplitudes (FTA), and raw time series (TS). PSD and EEG band features have been widely used in the literature in the past [43]–[45]. The PSD features are constructed from the periodogram or the power distribution of EEG signal over a series of narrow frequency bands, such as the frequencies from 1 Hz to the Nyquist frequency at 1 Hz steps, individually or jointly over all EEG channels [46], [47]. We used PSD constructed over the frequency intervals of 1 Hz from 1 Hz to 100 Hz (the Nyquist frequency). The PSD was calculated individually for each EEG channels separately and all PSD values were combined into a single feature vector of length *N_f_* = 21×100 = 2100 features.

The EEG band features are calculated as the EEG signal’s power in standard EEG bands defined conventionally as delta (1-4 Hz), theta (4-8 Hz), alpha (8-12 Hz), beta (12-30 Hz), and gamma (30-50 Hz) [48]. We calculated the EEG band-power values for each EEG channel separately and combined those into a single feature vector of length *N_f_* = 21× 6 = 126 features.

FTA features are considered newly in this work. For these, we perform a discrete Fourier transform (DFT) on each data frame, 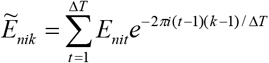, and use the DFT amplitudes directly in place of the features describing the EEG signal in each data frame for the BCI decoder. Whereas such FTA features are complex, machine learning algorithms require the features to be real. For that reason, we represent FTA features as real tuples in Cartesian, 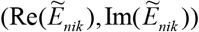, or polar, 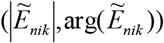, form. The total length of the FTA feature vector is N_*f*_ = 21× 200 = 4200 features.

The FTA feature vectors were additionally normalized to a zero total phase by means of the transformation 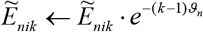, where 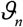 was an additional phase chosen so that 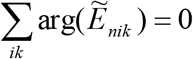 separately during each trial. This was done to remove random phase variation in FTA features due to minor changes in participants’ reaction time. It is well known that EEG BCI can suffer from variation in the initial point of ERP response curves such as related to the trial-to-trial variation in participant’s reaction time. This offset-time variation contributes a random phase shift to the DFT of each trial. While PSD and EEG band features are insensitive to such variation, because these features are defined as the square of the Fourier transform amplitude (from which the overall phase cancels out), such phase jitter can affect FTA features and the learning of motor imagery decoder based on the FTA feature vectors. To avoid this problem, we normalize the FTA feature vectors to have a zero total phase which removes such overall phase dependence.

After calculating the feature vectors for each trial, SVM and LDA machine learning algorithms were used to build decoders classifying each data frame as a specific mental imagery. We used the implementation of these algorithms in the Statistics Toolbox of Matlab. For SVM, we implemented a multiclass classifier following 1-vs-1 voting scheme and using the Matlab Statistics Toolbox’s “svmtrain” function. For LDA, we directly used the Matlab Statistics Toolbox’s “classify” function modified to allow application in the situation when the number of available examples was smaller than the number of features, such as present here. SVM and LDA methods are the examples of linear machine classification known for their strong performance and discriminative power in very high-dimensional and small dataset situations. LDA and SVM both have been used widely and enjoyed a significant success in EEG BCI as well as many other applications, for example see [49]–[52]. We do not go further into the details of these established algorithms here, whereas extensive literature is available on the subject for interested reader, see [53]–[58].

The classifiers were trained and their performance was evaluated using the cross-validation method, standard in the machine learning literature, using either randomized and sequential hold-out cross-validation with 70% of data used for training and 30% used for validation purposes.

### C. Online data processing

For interactive BCI control, we implemented the EEG data analysis system and an interactive real-time BCI application using Matlab. The interactive graphical user interface (iGUI) that we developed acquired raw EEG data from EEG-1200 in real time and performed learning of the BCI decoder as well as applied such decoder to offer a real-time control of a 3D robot manipulator arm simulated in on computer screen as shown in Fig. 2.

During the training sessions of the online experiments, the mentioned application collected a certain number of EEG data frames exemplifying different user’s motor imageries, as instructed by a pseudorandom training program. Typically, 300 motor imagery examples were collected during the training session and thus used to build the EEG BCI signal decoder, as described above. The data frame selection of [0,0.85] sec, FTA features in Cartesian form, and multi-class SVM were used for such a final decoder. After the training session, during the practice and the test sessions, the EEG data was acquired, data frames were selected and processed by the decoder online, in order to estimate the motor imagery class in real time and carry out the robot arm’s movement guided by the EEG signal according to a specific BCI control model to be described below.

In this work, two BCI control models were employed, a 3-state control model and a 6-state control model. In the 6-state control model, each of the five distinct motor imageries and one passive state were used as a control signal for the simulated robot arm. Each motor imagery was bound to one type of motion of the robot arm: Left and right hand movement imageries were bound to the motion of robot arm in left and right direction; leg movement imageries were bound to the forward and backward motion of the robot arm, tongue movement imagery was bound to the arm’s grabber hold or release motion, right panel in Fig. 3. The passive state was used to continue a previously started movement. Each motor imagery was used to initiate one motion in corresponding direction, and the motion continued until a second presentation of the same imagery or a presentation of a different motor imagery was encountered. The second observation of the same motor imagery stopped the motion of the robot arm while the observation of a different motor imagery stopped the current motion and initiated the new motion of the arm.

**Fig. 3.**
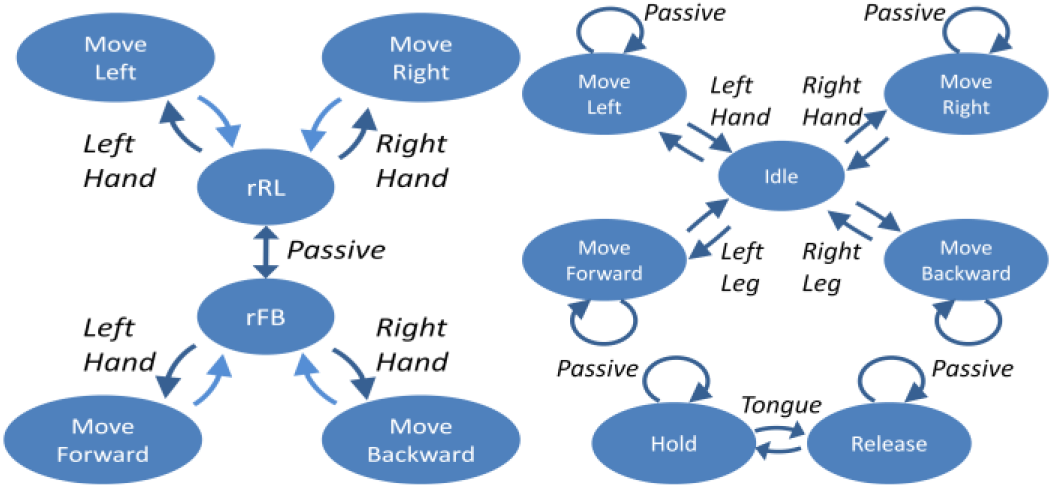
(Left) Schematic representation of the 3-state BCI control model. Left and right hand movements are used to move the manipulator either left and right or back and forth depending on motion regime switched by passive imagery implementation. (Right) Schematic representation of the 6-state BCI control model. Each motor imagery initiates one type of motion of the robot arm that continues through passive imagery presentation until a second presentation of the same or a new presentation of another motor imagery.

**Fig. 4.**
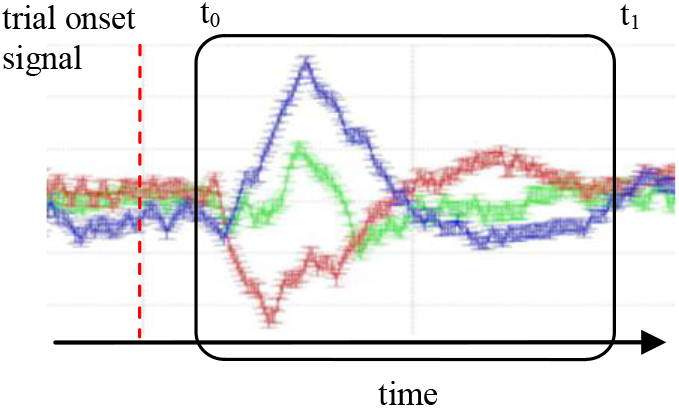
Illustration of the trial onset-locked decoding data frame used to detect different mental imagery states in the EEG BCI signal.

In the 3-state control model, only left-right hand and passive imageries were used. Left and right hand movement imageries were used to move the manipulator either left and right or back and forth, depending on the regime that could be switched among by the user using the passive imagery, left panel in Fig. 3. For example, a user could move the manipulator left or right by implementing left or right hand movements first, then, in order to move the manipulator forward or backward, the user could stay passive for one round during which the BCI would switch from left-right to forward-backward motion regime, and then continue using the left or right hand movements to move the manipulator forward or backward, respectively.

In order to switch the regime back, the user again could remain passive for one period. The observation of hand movement imageries moved the robot one step in the corresponding direction. Therefore, continuously moving the robot arm required the user to implement the desired imagery continuously over a sequence of “on”-signal presentation periods.

Our offline experiments indicated that both control strategies in principle could provide a similar average information throughput rate. However, we found in practice that substantially higher error rate of the 6-state control model proved to be frustrating for BCI users. Specifically, in our experiments the detection of the 3 mental imageries employed in the 3-state control model could be performed on average with 80-90% accuracy (see Results). At the same time, the detection of 6 mental imageries employed in the 6-state control model could be done with 50-60% accuracy. While these indicate that either 3-or 6-state control model provide similar information throughput of 1.2-1.5 bit per trial (0.4-0.5 bps), in practice, experiencing 1 in 2 error rate while attempting to control the 6-state BCI proved extremely frustrating and demotivating for the participants, negatively affecting their performance and resolve to master the control of the BCI.

For this reasons we focused on the 3-state BCI control model in our experiments.

## III. Results

### A. Discrimination of different mental imageries in offline EEG BCI

Twelve volunteers participated in this study after giving written informed consent. All participants were healthy individuals in their twenties, selected among the students of the Faculty of Engineering at Toros University and the Department of Physics and the Department of Biophysics at Mersin University, Mersin, Turkey. All participants took orientation session informing them about the experiments’ purpose, procedures, and their rights with respect to the collected information. All experimental procedures of the study had been reviewed and approved by the ethics committees of Toros University and Mersin University, Mersin, Turkey.

In all experiments, we observed that the mental imagery states could be inferred from the EEG data with accuracy significantly above chance. In the case of two motor imageries consisting of left and right hand movements, 6 out of 12 participants (50%) demonstrated the discrimination of left and right hand movement imageries from the EEG signal at above 95% level, and 8 out of 12 participants (67%) showed the discrimination of the two imageries at or above 90% level. This performance was stable across the experiments, with the top 6 individuals consistently demonstrating the discrimination accuracies better than 90% in all experiments. At the same time, for 2 out of 12 individuals (17%) a satisfactory performance of the BCI could not be achieved, showing at best 70-80% accuracy on these two imageries. Both SVM and LDA-based decoders showed similar results, with SVM showing a marginally better performance, on average outperforming LDA by 3-5%. These results are summarized in Table I. The discrimination of two mental imagery states can be used for binary yes/no communication with BCI and similar decisions.

**TABLE I.** Participants performance in 2-state BCI discrimination task

| Participant | Performance SVM |  |  | Performance LDA |  |  | Number of experiments |
| --- | --- | --- | --- | --- | --- | --- | --- |
|  | Average | Worst | Best | Average | Worst | Best |  |
| BA | 0.77 | 0.75 | 0.79 | 0.73 | 0.69 | 0.77 | 2 |
| EM | 0.96 | 0.93 | 1.00 | 0.95 | 0.92 | 0.97 | 3 |
| ER | 0.95 | 0.89 | 0.99 | 0.94 | 0.88 | 0.97 | 3 |
| HI | 0.98 | 0.96 | 0.99 | 0.94 | 0.88 | 0.98 | 5 |
| ME | 0.75 | 0.66 | 0.84 | 0.70 | 0.62 | 0.77 | 2 |
| MR | 0.88 | 0.74 | 0.92 | 0.79 | 0.73 | 0.86 | 5 |
| SE | 0.95 | 0.89 | 1.00 | 0.92 | 0.82 | 0.98 | 5 |
| YU | 0.91 | 0.78 | 0.93 | 0.84 | 0.72 | 0.92 | 5 |
| EK | 0.84 | 0.76 | 0.93 | 0.85 | 0.79 | 0.90 | 2 |
| ES | 0.99 | 0.98 | 0.99 | 0.98 | 0.97 | 0.99 | 2 |
| UL | 0.81 | 0.76 | 0.87 | 0.79 | 0.74 | 0.84 | 2 |
| YL | 0.98 | 0.98 | 0.98 | 0.97 | 0.96 | 0.97 | 2 |
| Overall | 0.90 | 0.66 | 1.00 | 0.87 | 0.62 | 0.99 | 38 |

While such mode of operations could be realized by our BCI with high efficiency, the binary communication paradigm is too limited for control of a robot manipulator system of any meaningful complexity. For that reason, we considered the addition of a 3^rd^ mental imagery state to our BCI. The 3^rd^ state can be used to alternate between different operating regimes of a robotic manipulator system, for example, such as left-and-right and forward-and-backward motion, whereas the original two mental imageries can be used to move the manipulator in the two directions of each of those regimes. We choose the “passive” mental imagery state as such a 3^rd^ state addition. The passive state is a natural choice for this purpose, whereas the user only needs to remain passive - that is, don’t do anything intentionally, in order for the BCI to switch operating regimes. This choice is clearly advantageous to the choice where a conscious action would be required to switch BCI regime, whereas the user would be forced to remain active continuously as the BCI switches regime even as no net action was produced.

With respect to the 3-state BCI communication paradigm, in our experiments we observe that the detection of respective mental imageries can still be performed with high accuracy. For 5 out of 12 participants (42%) we observe that the 3 mental states can be discriminated at better than 85% accuracy. The second group of 5 best individuals (42%) shows significant but worse performance of 70-80%, and 2 out of 9 participants (22%) do not achieve satisfactory performance, with the accuracy of detection of these states between 40 and 60%, Table II.

**TABLE II.** Participants performance in 3-state BCI discrimination task

| Participant | Performance SVM |  |  | Performance LDA |  |  |
| --- | --- | --- | --- | --- | --- | --- |
|  | Average | Worst | Best | Average | Worst | Best |
| BA | 0.54 | 0.49 | 0.58 | 0.55 | 0.47 | 0.62 |
| EM | 0.85 | 0.80 | 0.89 | 0.82 | 0.81 | 0.83 |
| ER | 0.89 | 0.83 | 0.93 | 0.84 | 0.74 | 0.89 |
| HI | 0.89 | 0.86 | 0.95 | 0.90 | 0.85 | 0.95 |
| ME | 0.52 | 0.42 | 0.62 | 0.52 | 0.41 | 0.63 |
| MR | 0.70 | 0.59 | 0.82 | 0.67 | 0.55 | 0.77 |
| SE | 0.82 | 0.63 | 0.90 | 0.78 | 0.60 | 0.88 |
| YU | 0.76 | 0.71 | 0.83 | 0.74 | 0.68 | 0.79 |
| EK | 0.71 | 0.61 | 0.81 | 0.70 | 0.58 | 0.81 |
| ES | 0.98 | 0.96 | 0.99 | 0.97 | 0.95 | 0.97 |
| UL | 0.69 | 0.58 | 0.80 | 0.66 | 0.57 | 0.75 |
| YL | 0.90 | 0.89 | 0.91 | 0.88 | 0.87 | 0.88 |
| Overall | 0.77 | 0.49 | 0.99 | 0.75 | 0.41 | 0.97 |

We now consider the task of detecting up to 6 different states in an EEG BCI. A 6-state BCI control paradigm can offer a significantly richer dynamics to users. In that setting, a control model can be designed where each mental imagery is mapped to a different motion of a 3 dof manipulator, including left and right rotation, forward and backward motion, and grabbing and releasing objects. The set of states used for the 6-state BCI communication model here are left and right hand movement imagery, left and right leg movement imagery, tongue movement imagery, and a passive imagery.

We observe a significantly above chance performance in all our experiments in this situation as well. However, the error rates became more significant. Specifically, top 5 out of 12 participants were able to demonstrate 75-90% accuracy in this BCI task as well, Table III, as compared to the baseline or “chance” performance of only 17%. These individuals’ performance is far above chance and may be interesting for further investigation. At the same time, the performance of the other individuals degraded significantly to 50-60% accuracy. While still significantly above chance, such rate of errors presents a serious obstacle on the way of utilizing this EEG BCI communication paradigm for practical purposes.

**TABLE III.**
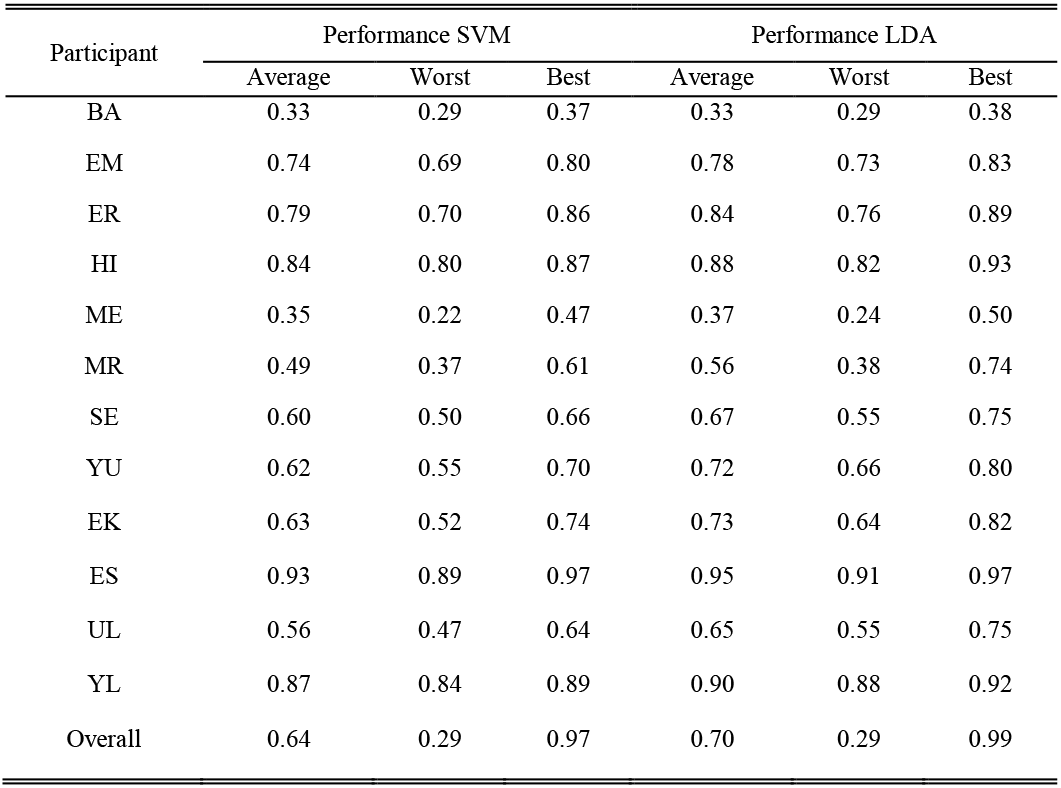
Participants performance in 6-state BCI discrimination task

| Participant | Performance SVM |  |  | Performance LDA |  |  |
| --- | --- | --- | --- | --- | --- | --- |
|  | Average | Worst | Best | Average | Worst | Best |
| BA | 0.33 | 0.29 | 0.37 | 0.33 | 0.29 | 0.38 |
| EM | 0.74 | 0.69 | 0.80 | 0.78 | 0.73 | 0.83 |
| ER | 0.79 | 0.70 | 0.86 | 0.84 | 0.76 | 0.89 |
| HI | 0.84 | 0.80 | 0.87 | 0.88 | 0.82 | 0.93 |
| ME | 0.35 | 0.22 | 0.47 | 0.37 | 0.24 | 0.50 |
| MR | 0.49 | 0.37 | 0.61 | 0.56 | 0.38 | 0.74 |
| SE | 0.60 | 0.50 | 0.66 | 0.67 | 0.55 | 0.75 |
| YU | 0.62 | 0.55 | 0.70 | 0.72 | 0.66 | 0.80 |
| EK | 0.63 | 0.52 | 0.74 | 0.73 | 0.64 | 0.82 |
| ES | 0.93 | 0.89 | 0.97 | 0.95 | 0.91 | 0.97 |
| UL | 0.56 | 0.47 | 0.64 | 0.65 | 0.55 | 0.75 |
| YL | 0.87 | 0.84 | 0.89 | 0.90 | 0.88 | 0.92 |
| Overall | 0.64 | 0.29 | 0.97 | 0.70 | 0.29 | 0.99 |

### B. Optimization of BCI decoder

In this section we consider several optimizations of BCI decoder parameters and design choices.

We first consider the optimization of the decoder’s data frame selection. In order to detect different mental imagery states in EEG BCI data, a trial onset-locked data frames, [*t_0_,t_1_*], are used for decoding as illustrated in Fig. 4. Selection of the parameters of such data frames is important for optimizing the discrimination ability of the BCI decoder. In particular, making the decoding frame too large can obscure the relevant features due to accumulation of irrelevant noise contributed by uninformative EEG signal fragments immediately prior and immediately thereafter the useful EEG BCI signal. Likewise, choosing the data frame that is too small can miss such relevant signal all together.

To select the best such data frame parameters, we scanned over all possible choices of the initial frame offset, *t_0_*, in the range from -0.5 sec to +1.0 sec (relative to trial onset signal) at a 0.1 sec increment. Likewise, we inspected different possible decoding data frame-length values, *dt=t_1_-t_0_*, in the range between 0.2 and 2.0 sec, again at a 0.1 sec increments. We performed a complete search on the grid of such points for best parameters for each participant and each experiment. Fig. 5 shows the examples of the 2-state BCI discrimination accuracies for one high-performing individual (HI), one intermediately performing individual (YU), and one low-performing individual (BA). Several features can be pointed out in that figure. First, at the top-left corner of the diagrams a region with chance performance of just about 50% is observed. Upon closer examination, this region corresponds to the data frames such that lie entirely prior to the trial onset time. As the information about the mental imagery cannot possibly be contained in the decoding frames lying entirely prior to the trial onset signal, the decoder performance in that part of the diagram is at the simple chance level.

The decoder’s performance increases sharply once the decoding frame begins to overlap with the region past the trial onset time at approximately 300-500 milliseconds. Indeed, the highest accuracy is consistently observed in the triangular region of the diagram defined by the frame initial offset *t_0_* of - 0.5 to 0.3 sec and the frame length 0.2 to 0.9 sec. A rapid drop in performance is observed when *t_0_* exceeds 0.5 sec limit. This indicates that the information related to the identification of mental imagery state is not present in the EEG signal after approximately 500 millisecond past trial’s onset signal. Finally, we observe similar structure of the *t_0_ -dt* diagrams in Fig. 5 for each participants, whereas the lower-performing individuals only exhibit a lower increase in decoder’s accuracy in the above mentioned best-detection triangle but otherwise the structure of the diagram remains the same.

**Fig. 5.**
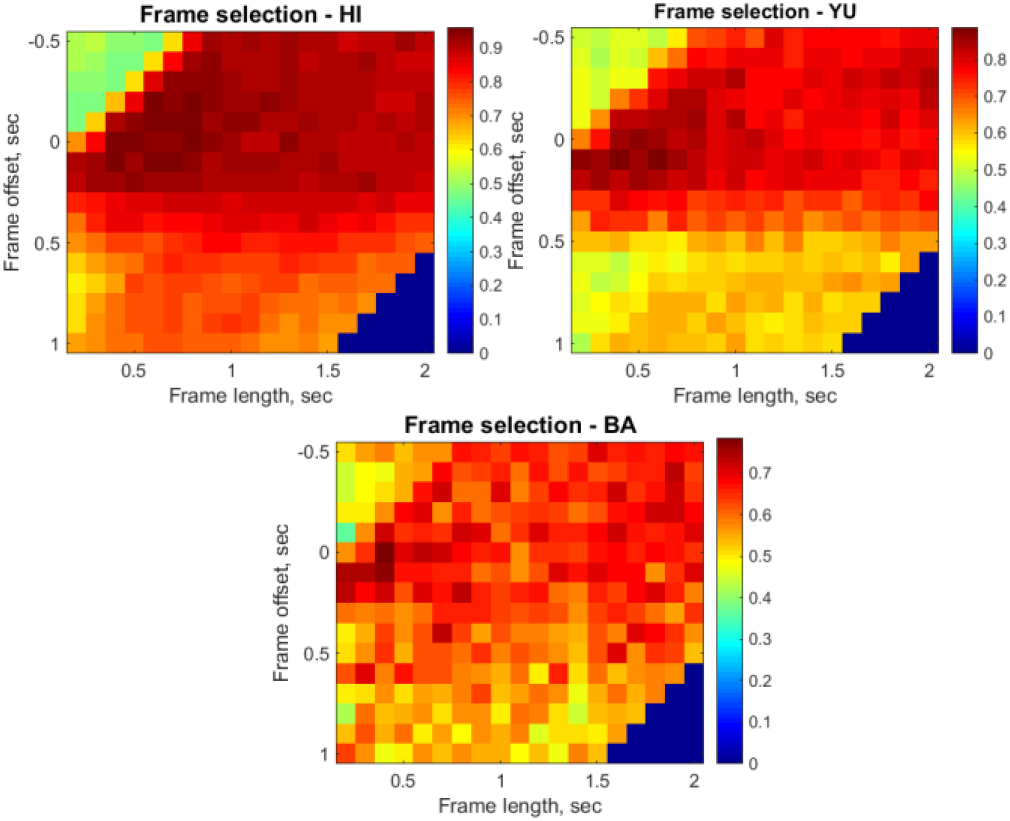
Diagrams showing the accuracy of 2-state BCI discrimination versus the parameters of the decoder’s data frame selection for one high-performing individual (HI), one intermediately performing individual (YU), and one low-performing individual (BA).

All participants demonstrate close to optimal performance near the choice of the decoding frame parameters *t_0_=0-0.2* sec and dt=0.6-0.9 sec. Thus, it appears possible to adopt a uniform choice of the decoding frame parameters [*t_0_,t_1_*]=[0,0.85] sec as a generally suitable for most participants, although fine-tuning of the decoding frame’s position can allow moderate increase in BCI decoder’s performance on individual experiment basis.

We second consider the issue of selecting the feature representation of EEG signal. The selection of feature representation of EEG signal for BCI decoder can significantly affect the BCI’s performance. We investigate this effect here by examine the performance of BCI decoders built using EEG band power features, power spectral density (PSD) features, as well as new Fourier transform amplitude features. For all the signal power-based features such as EEG band powers and PSD, we inspect the row power and the log10 (that is, the Decibel) representation of such features. The row time-series (TS) features are likewise considered in this section: The time-series features had been used in EEG BCI in some past works, but did not find widespread use in EEG BCI literature.

To briefly recap the above features’ definitions, if the amplitudes of the Fourier transform of the EEG signal within a certain signal frame [ř_0_, t] are a_ŕ_ (*f*), where *f* is the DFT frequency index and *i* is the index of EEG channel, then EEG band power features are defined as

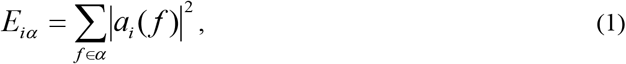

where α represent the frequency range of a standard EEG band such as alpha, beta, gamma, delta, or theta. PSD features are defined as

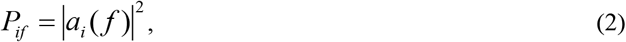

FTA features in polar form are defined as

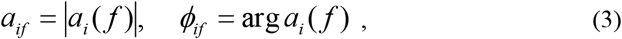

and FTA features in Cartesian form are defined as

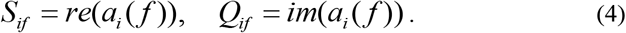

Note that FTA features are real-valued tuples for a range of frequencies *f* from 0 to *F_Nyquist_* = *F_Sample_* /2, while PSD is an array of single values for the same frequencies. In the decibel representation, EEG band power and PSD features are calculated as *H_iα_* = 10log_10_ *E_iα_* and *D_if_ = 10log_10_ P_if_*.

We note that EEG band power, PST, and polar-form FTA features form a hierarchy of feature sets of progressively increasing information content. While EEG band power features describe only 5 to 6 different values providing the power of the EEG signal in standard EEG bands, PSD features provide up to 100 such values, for example, for every 1 Hz bands in a 1-100 Hz frequency range. While EEG band power and PSD features lack the information about the phase of the Fourier transform corresponding to the specific shape of the EEG waveform observed in a particular BCI trial, that information is now recovered in the FTA-polar feature set, which therefore can be seen as an equivalent to the square root of PSD with an addition of the phase information over all frequencies. FTA-polar is the first feature set containing the complete information about the EEG waveform and allowing the complete reconstruction of the original EEG waveform. The FTA-Cartesian feature set is an equivalent of the FTA-polar, since both are produced from the complex DFT amplitudes. However, the relationship between the two is nonlinear and, therefore, the performance of a simple linear machine classifier such as SVM or LDA cannot be expected to be the same on the FTA-polar and FTA-Cartesian feature sets. In fact, we observe the same (see below). Finally, FTA-Cartesian features are a linear combination of the EEG time-series features (TS), whereas *S* and *Q* are simply the real and imaginary parts of the Fourier transform viewed as a filter applied to the TS data. The performance of linear machine learning classifiers can be expected to be similar on these two feature sets.

We test the performance of LDA and SVM-based BCI decoders using all of the above described feature sets, Table IV. While we observe that performance remains well above chance for all the above choices of EEG signal representation, FTA-Cartesian features perform dramatically better than all of the other feature sets including EEG band power, PSD, and FTA-polar. Whereas those choices yield an all-subject-average performance of close to 80% on the 2-state BCI discrimination task, FTA-Cartesian features yield average performance close to 90%. For 3-state BCI, the performance achieved using the power-based features is 55-60%, whereas FTA-Cartesian achieved yields close to 75% BCI state discrimination accuracy. In the BCI task involving 6 mental imagery states, the average performance achieved with EEG band power, PSD, and FTA-polar feature sets is 40% versus 65% for FTA-Cartesian. Among the power-based features and their decibel representations, we observe no significant differences in the decoder’s performance. Therefore, FTA-Cartesian features are observed to be far superior to the choice of signal power-based EEG features such as EEG band power or PSD, commonly employed in the literature.

**TABLE IV.** Performance of different feature representations for mental imagery state discrimination task

|  | EEG bands | log-EEG bands | PSD | log-PSD | FTA Polar | FTA Cartesian | Time series |
| --- | --- | --- | --- | --- | --- | --- | --- |
| SVM-2 | 0.770 | 0.769 | 0.780 | 0.766 | 0.812 | 0.910 | 0.897 |
| SVM-3 | 0.579 | 0.594 | 0.580 | 0.560 | 0.624 | 0.744 | 0.757 |
| SVM-6 | 0.439 | 0.427 | 0.422 | 0.384 | 0.432 | 0.594 | 0.597 |
| LDA-2 | 0.698 | 0.710 | 0.740 | 0.717 | 0.769 | 0.860 | 0.858 |
| LDA-3 | 0.578 | 0.612 | 0.586 | 0.563 | 0.622 | 0.754 | 0.734 |
| LDA-6 | 0.443 | 0.439 | 0.453 | 0.415 | 0.493 | 0.646 | 0.605 |

Since FTA-Cartesian features are a simple linear transformation of the EEG time-series, we can expect that the linear machine learning classifiers such as SVM and LDA will produce comparable performance on these feature sets. Indeed, we observe that in our experiments: both FTA-Cartesian and TS features provide similar performance for discrimination of up to 6 mental imagery states. However, as we will see below, FTA features have the added advantage of a more concise representation of the EEG signal with respect to frequency decomposition, which allows for a far smaller subset of FTA features to be used in the EEG BCI decoder without sacrificing the performance. Therefore, we conclude that FTA-Cartesian features are the better choice for our EEG BCI.

Finally, we consider the issue of the referencing of the EEG data. When EEG data is acquired by an EEG acquisition device, it is recorded with respect to a particular voltage reference [48]. Choosing a different voltage reference affects the final representation of the EEG signal by adding or subtracting a time-varying common mode component. We inspect the impact of different choices of such referencing mode on the performance of EEG BCI decoders. Specifically, we examine the choices including the system’s 0 Volt reference (in the Nihon Kohden EEG-1200 system), A1-A2 average reference (defined as the average of the voltages on A1 and A2 electrodes), the common reference (defined as the average potential of all EEG electrodes), and the Laplace reference (defined as the average of 4 neighbor electrodes for each electrode). The results of this analysis are presented in Table V. overall, we observe that the choice of reference voltage can have a noticeable impact on the EEG BCI performance for EEG band powers and PSD features. A BCI state detection accuracy improves up to 5% consistently when such references are used versus the system 0 Volt. In the case of FTA-Cartesian and TS features, the differences that can be attributed to the change of voltage reference are much less pronounced, with Laplace reference showing marginally better results.

**TABLE V.**
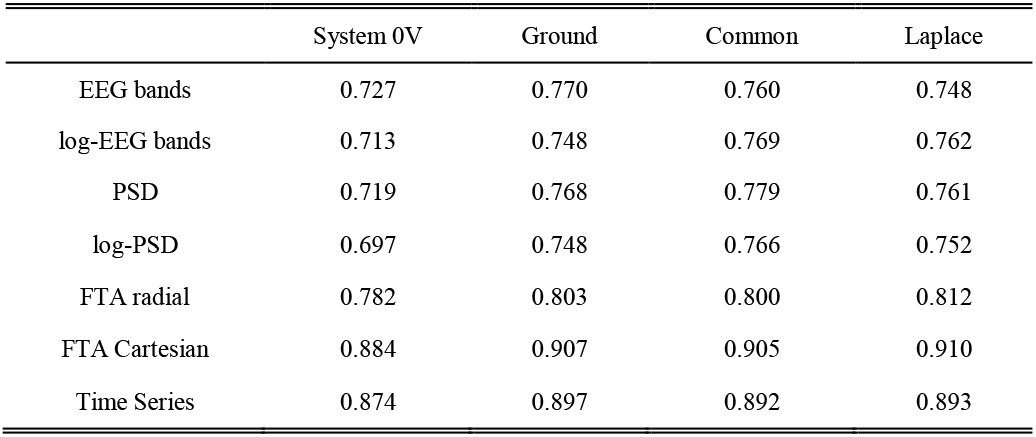
The impact of different choices of voltage reference on the BCI states discrimination accuracy

### C. The information content of EEG signal by frequency range

Preprocessing of EEG signal by using low-pass filters is a common practice in EEG BCI [48], [59]–[66]. In this work, we also observed that applying a low-pass filter to EEG signal can improve the performance of BCI decoder. In the case of the frequency-space features such as PSD or FTA, such a filtering can be implemented simply by rejecting those features that not fall into the pass-range [0, *F*c]. In the case of TS features, we apply low-pass filter using 8^th^-order Butterworth low-pass filter. In either case, the effect of discarding high frequencies from EEG signal is found to be advantageous for the performance of EEG BCI. In fact, our experimentation suggests that keeping only the lowest frequency ranges of 0 to 5-10 Hz can result in the best overall performance of EEG BCI.

The above experimentation suggests that most of useful BCI information in EEG signal is contained at very low frequencies. To investigate this issue further, we performed a series of numerical experiments in which the BCI decoder was constrained to use only the EEG data coming from a narrow 5 Hz-wide band, chosen in the frequency range 0-80 Hz.

In other words, we first inspected the BCI decoder using only the EEG data filtered to the frequency range of 0-5 Hz. Second, we inspected the BCI decoder using the EEG data filtered in the frequency band 5-10Hz, and so on. We considered the EEG BCI decoders for 2, 3, and 6 mental state discrimination. In all cases, we observed that the ability of EEG BCI decoders to discriminate mental imageries degraded dramatically as the higher-frequency bands were selected, reducing to essentially the chance level after the threshold of approximately 20 Hz, Fig. 6. We conclude that the information relevant to the task of discriminating different motor imageries in an EEG BCI resides primarily in the low frequency range of 0-15 Hz, with the most such information residing in the range 0-10 Hz.

**Fig. 6.**
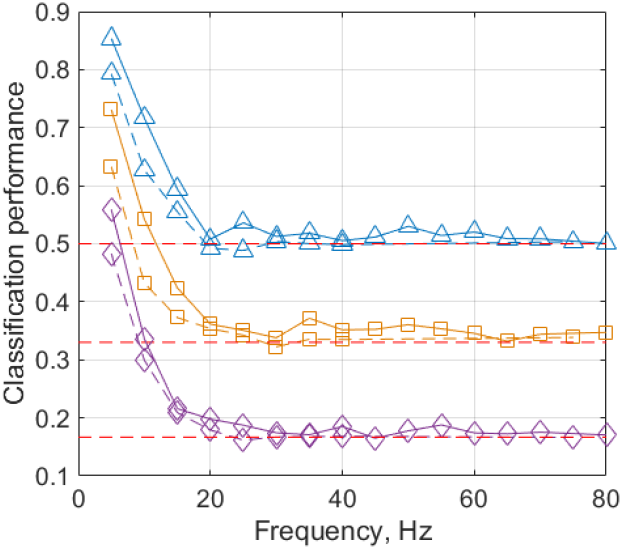
The performance of EEG BCI decoder constrained to use narrow 5 Hz frequency bands in the range 0-80 Hz. Solid lines - FTA-Cartesian features sorted by 5 Hz frequency bands from 0-5 Hz to 75-80 Hz. Dashed lines - TS features sorted by 5 Hz frequency bands. Triangles indicate 2-state BCI classification, squares indicate 3 state BCI classification, and diamonds are 6-state BCI classification. Red dashed lines indicate the chance level for 2, 3, and 6-state classification task. In all cases the classification performance is seen to drop to the chance levels at the frequency bands exceeding approximately 20 Hz.

### D. Consistency of BCI performance across participants

In our experiments, we observe that the same participants tended to perform better in all experiments and for all types of BCI tasks, while the other same participants always tended to perform worse in all experiments. Table VI illustrates this observation by pulling together the performance data of different participants over all inspected BCI tasks and all experiments. As can be seen in Table VI, the top performing individuals perform better in all BCI tasks and all experiments. Likewise, the intermediate-level individuals performed at an intermediate level in all tasks, while the low-performing individuals always showed low levels of performance.

**TABLE VI.** Consistency of BCI performance of different individuals across BCI tasks, ordered by the participants’ overall performance level

| Rank | Participant | Average performance |  |  |
| --- | --- | --- | --- | --- |
|  |  | 2-state | 3-state | 6-state |
| 1 | ES | 0.99 | 0.98 | 0.93 |
| 2 | YL | 0.98 | 0.90 | 0.87 |
| 3 | HI | 0.98 | 0.89 | 0.84 |
| 4 | ER | 0.95 | 0.89 | 0.79 |
| 5 | EM | 0.96 | 0.85 | 0.74 |
| 6 | SE | 0.95 | 0.82 | 0.60 |
| 7 | YU | 0.91 | 0.76 | 0.62 |
| 8 | EK | 0.84 | 0.71 | 0.63 |
| 9 | MR | 0.88 | 0.70 | 0.49 |
| 10 | UL | 0.81 | 0.69 | 0.56 |
| 11 | ME | 0.75 | 0.52 | 0.35 |
| 12 | BA | 0.77 | 0.54 | 0.33 |

### E. Online EEG BCI applications

We implemented an online, interactive EEG BCI system based on the hardware and the software described in the previous sections. The implemented system contained a virtual 3 dof-robot manipulator simulated in 3D on computer screen and capable of lateral motion, longitudinal motions, and hold-and-release motion. The manipulator could be controlled interactively by users by means of the EEG BCI, Fig. 7.

**Fig. 7.**
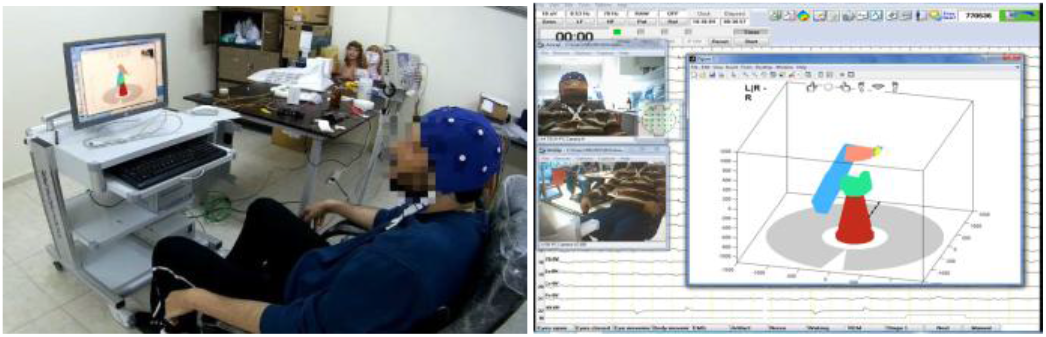
Online experiments in which the participants have controled a 3 dof-robot manipulator arm using EEG BCI.

Three best performing individuals were invited to participate in the interactive BCI trials. Each experiment consisted of three 15-minute sessions as described elsewhere in this work: a BCI decoder training session, a practice session, and a test session. During the training session, the users taught the BCI decoder in supervised manner by implementing a series of mental imageries as indicated to them by the GUI of the BCI. The robot manipulator did not move during that time and the BCI only accumulated the data to be used for training the BCI decoder and did not respond to the user. After the conclusion of training, the participants were invited to attempt to control the manipulator interactively, in a free-style exploration manner by using the trained decoder and a 3-state BCI control model (see Methods). Finally, in the test session the participants were given a series of task to move manipulator to different positions and attempted to complete them. The tasks included moving the manipulator in two (left-right or forward-backward) to four (left-right and forward-backward) directions for up to 4 steps. The participants completed in total 7-8 tasks during the 15-minute test sessions.

The participants’ performance was evaluated as the percentage of the tasks completed successfully, the average time taken to complete a task, and the average accuracy of the control of the manipulator. The control accuracy was quantified as the percentage of the manipulator moves that were in a correct directions moving the manipulator towards the target. The best participant (HI) had shown in the interactive trials the average control accuracy of 84.6%, and the second best participant (ER) had shown in the interactive trials the control accuracy of 77.8%, which was consistent with the results observed for these participants in the segment of this work’s analyses performed offline.

The third participant (ES) experienced greater difficulties controlling the BCI, being able to control the BCI with an average accuracy of only 49.6%, far below the performance levels shown in the offline experiments. The successful participants required 7 to 10 seconds to implement one manipulator move, on average. The participant HI spent 6.5 seconds and the participant ER required 9.3 seconds to implement each manipulator’s move. Due to the BCI control model here, the BCI could accept on average one command per a 3 second period, equal to the time per one “on”-signal’s presentation in the BCI. Together with the time necessary for switching the regime of motion of the BCI, this implied on average 4.1 seconds per manipulator move - the best case scenario. The best participant, therefore, was able to control the manipulator with 50% time overhead and 15% error rate, while the second best participant required approximately twice the ideal amount of time and made close to 25% errors controlling the BCI. Due to difficulty controlling the BCI that the third participant experienced, that participant spent from 13 to 40 seconds per move in successful trials, being able to complete only 40% of assigned tests. The other two subjects were able to complete 100% of the tasks given to them. The results of interactive BCI application are summarized in Table VII.

**TABLE VII.**
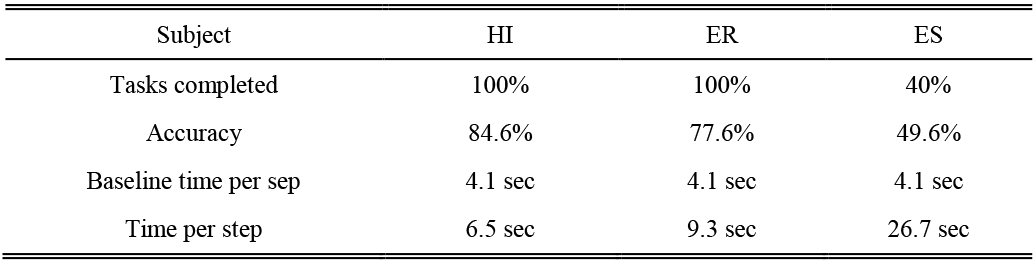
Participants’ performance in online BCI experiments

## IV. Discussion

In our experiments, we distinguish as many as 6 mental imagery states in EEG BCI well above chance level. The observed accuracy of discriminating mental imagery states achieved here is 90% for 2 mental imageries (right and left hand motor imageries), 77% for 3 mental imageries, and 64% for 6 imageries, averaged over all subjects. Our results for offline experiments are consistent with those found in the literature [67]–[73]. In [67] accuracies of discriminating two mental tasks for 3 subjects have been found as 94%, 90% and 84%, respectively. In [68] classification accuracies for 3 subjects of 86%, 96%, and 90% for two mental tasks, and 46%, 67%, and 45% for up to five mental tasks have been reported using EEG signal band power features and a hidden Markov model for electroencephalographic patterns classification. In [69], experimental EEG recordings have been obtained for 3 subjects while performing 3 different metal task and BCI discrimination accuracies of up to 62% have been reported. In [71], the average of classification accuracies of six subjects of up to 75.6% have been reported using 2 mental imageries (right and left hand finger motor imageries). In [70], 18 participants participating in experiments aimed at learning control of EEG BCI by using 3 mental imagery tasks have shown mean group performance of 53%, consistent with our results. In [72], Echo State Networks have been used for classification of EEG signals in mental task-based brain-computer interface. In this study, classification accuracies as high as 95% have been reported using two mental task and up to 65% using four mental tasks. The group averages of 81% for two-task classification and 9 participants, and 54% for four-task classification have been reported. In [73], for a group of 5 subjects the classification accuracies of four mental imageries (opening and closing of left and right hand, two hands, and feet movements) of 86.6 *%* have been reported. The group-average performances of 90% for two tasks, 77% for three tasks, and 64% for six tasks reported in this work, therefore, are consistent and in many cases superior to similar results reported in the literature.

We study the impact of different choices in the design of EEG BCI decoder parameters on BCI performance, including the choice of EEG signal’s feature representation, the choice of decoder’s detection frame parameters, the choice of voltage reference, and other choices. We observe that FTA features show superior performance outperforming by close to 30% all of the features based on calculating the EEG signal power spectrum, such as used in the EEG BCI literature conventionally. In this work, the EEG signal power features are shown to achieve up to 78% discrimination accuracy on 2-state BCI discrimination task, average over all participants, whereas FTA features allow the same discrimination to be performed with greater than 90% accuracy. For discrimination of 6 mental imageries, the EEG signal power features allow up to 45% discrimination accuracy, while the use of FTA features results in 65% accuracy on average over all participants and up to 85-90% for best individuals. The use of FTA features in this work, therefore, allows achieving a significant improvement in the performance of EEG BCI.

Low-pass filtering of EEG signal prior to its processing in BCI is a common practice in EEG BCI literature. We also observe that low-pass filtering can help improve the performance of the decoder in the EEG BCI. Furthermore, we observe that low-pass filtering the EEG signal to only 5 Hz high-cutoff results in some of the best performance of the BCI. This suggests that most useful information about the motor-based mental imagery used in this work resides at very low frequencies in the EEG signal.

Indeed, similar results - that the useful information in EEG BCI may be encoded by the lowest EEG frequency bands - is also indicated in past studies including such of upper limb movement intentions [74]–[77], studies involving field potential [78]–[81], ECoG [29], [30], [82], and closed-loop studies with implanted intra-cortical electrodes [83]. On the other hand, traditionally motor-imagery EEG BCI have used power modulation in higher frequency bands of EEG signal such as mu (8-13 Hz) and beta (20-30 Hz) rhythms.

We inspect this situation in greater depth by testing the discriminability of mental motor imageries using the EEG data filtered to different narrow 5 Hz-bands ranging from 0-5 Hz to 75-80 Hz. We observe that the ability of EEG BCI to distinguish such motor imagery states drops rapidly as the frequency of the EEG filtering band increases past 15-20 Hz. Only 0-5 Hz, 5-10 Hz and 10-15 Hz frequency bands allow the EEG BCI performance that is significantly different from chance, and only the frequencies of 0-5 Hz and 5-10 Hz show such performance better than the chance by a meaningful margin. Specifically, in the case of discriminating the motor imageries of left and right hand movements, the all-participants average discrimination accuracy observed using the EEG signal in 0-5 Hz band in our experiments was 80-90%, for 5-10 Hz - 65-75%, and for 10-15 Hz - 55-60% - just slightly above the chance level of 50%. Beyond 15 Hz, the discrimination accuracy for this BCI task was not different from chance. At the same time, for discriminating 6 mental imagery states, the all-participants average accuracy was 50-60% when using 0-5 Hz frequency band, 30-35% when using 5-10 Hz band, and 20-23% for 10-15 Hz band, with the higher bands again showing no difference from the chance level of 16%. We conclude that the low frequencies of 0-15 Hz carry the most significant information related to the different motor imageries used in EEG BCI, with most of such information coming from the frequency band of only 0-10 Hz.

In our study, the participants demonstrated consistent performance when grouped into high-performing, intermediate-performing and low-performing groups. 5 out of 12 participants demonstrated high level of performance using our BCI, achieving in the offline experiments the accuracy on 2-state BCI task of 95-100%, 3-state BCI task of 85-90%, and 6-state BCI task of 80-90%. These individuals demonstrated high performance related to all BCI tasks and consistently in all experiments. In online BCI trials, these participants showed good ability to control the BCI in interactive settings demonstrating the ability to execute motions using a simulated 3D robotic manipulator arm and the BCI with 80% accuracy at a rate of 6-9 moves per minute. The ability of these individuals to control the BCI presents interest for further practical application. 5 out of 12 participants demonstrated intermediate levels of performance achieving the accuracy of 80-90% in 2-state BCI task, 70-80% in 3-state BCI task, and 50-70% in 6-state BCI task. 2 out of 12 participants could not achieve a satisfactory BCI control ability. While these individuals still showed performance significantly above chance while using our BCI, they could achieve the BCI control accuracy of only 75% in 2-state scenario, 50% in 3-state scenario, and 35% in 6-state scenario.

In our online experiments, a single 15-minute session was used for training the BCI decoder. This was done because the participants’ intent during the practice and the test sessions deemed to not be reliably known, given the design choices made in our experiments. In principle, such design can be seen as suboptimal, whereas suggestions for decoder learning involving also the practice and the test sessions have been made in the literature [84], [85]. One common approach for that can be to treat the BCI moves consistent with the target pursued by the user in a test or practice session as correct. on the other hand, other studies have suggested that learning BCI can be viewed as a process similar to learning the use of any other tool [22]. Thus, the process of BCI application can be structured intentionally in such a way that a moderately successful BCI decoder is offered to the users first, and then the users adapt to the decoder, which happens by changing of the users’ neural patterns in the brain, to better grasp the control of the fixed BCI. In this scenario, the BCI decoder needs to remain fixed intentionally, in order to assist the user in that adaptation process [18]. Distinguishing among these two possibilities is one of the future targets for our work.

In EEG BCI, high session-to-session variability of the EEG signal and accompanying need for re-training the BCI is well known [86]–[89]. While the reasons for that variability are not well understood, they may include session-to-session variability in electrode placement, changes in scalp and electrode impedance due to humidity, temperature, channel motion, sweat, as well as emotional, hormonal, and pharmacological changes. Similarly, in our experiments we observe the need for retraining or re-calibrating the BCI decoder in each experiment. The necessity to train the decoder for each new application of the BCI is a disadvantage of existing EEG BCI technology. However, in this work the training session lasted for only 15 minutes, and after that the same decoder was used until the end of the experiment. This training time is much shorter than that described in other BCI studies [58], [90]–[92].

## V. Conclusions

In this work, we study the possibility of implementing an EEG-based BCI for high-performance control of an assistive robot manipulator in 3 dimensions. For this purpose, we carry out an extensive offline study of mental imagery discrimination using EEG BCI as well as perform online BCI control experiments. We use motor imagery BCI control paradigm successful in the literature. We inspect different design choices of such an EEG BCI. Our results indicate that the levels of performance interesting for practical applications can be achieved using EEG BCI with only noninvasive neural activity imaging modality. The option for providing the BCI control noninvasively is of great interest for BCI development. In that respect, EEG is of special interest because of its low cost, technological maturity, robustness, and the ease of use with modern devices. In our experiments, best individuals demonstrated up to 80-90% control accuracy with up to 6 BCI states in offline settings, and 80-90% control accuracy using 3 mental imagery states in interactive, online settings.

Despite these achievements, it appears that incorporating relatively high levels of control errors into EEG BCI systems with any practical applications is eventual inevitability. The EEG signal is well known to be affected by highly significant uncontrollable variations due to a variety of factors including noise as well as systematic variations such as subjects’ emotional and mental condition and external factors such as the condition of skin and environment. This makes achieving faultless extraction of BCI intent from EEG data implausible. The future EEG BCI systems, therefore, will need to accommodate for this eventuality.

Another interesting observation is that different individuals appear to demonstrate distinctly different abilities in controlling the EEG BCI. In our study, a group of several individuals demonstrated consistently high BCI performance, while a group of other individuals was not able to achieve satisfactory performance at all. This observation is consistent with the notion of “BCI literacy” in modern EEG BCI literature [47], [93], [94]. Certain inherent, either psychological or physiological parameters may be responsible for this variability in the ability of different individuals to control EEG BCI. If so, focusing on the design of EEG BCI tailored to specific individuals with the ability to control them may be a plausible strategy for the development of practically applicable EEG BCI.

